# Joint effect of changing selection and demography on the site frequency spectrum

**DOI:** 10.1101/2022.02.19.481143

**Authors:** Kavita Jain, Sachin Kaushik

## Abstract

The site frequency spectrum (SFS) is an important statistic that summarizes the molecular variation in a population, and used to estimate population-genetic parameters and detect natural selection. While the equilibrium SFS in a constant environment is quite well studied, recent research has focused on nonequilibrium SFS to elucidate the role of demography when selection is constant in time and of fluctuating selection in a population of constant size. However, the joint effect of time-dependent selection and population size has not been investigated so far. Here, we study the SFS in a randomly mating, diploid population in which both the population size and selection coefficient vary periodically with time using a diffusion theory approach, and derive simple analytical expressions for the time-averaged SFS in slowly and rapidly changing environments. We show that for strong selection and in slowly changing environments, the time-averaged SFS differs significantly from the equilibrium SFS when the population experiences both positive and negative cycles of the selection coefficient. The deviation depends on the time spent by the population in the deleterious part of the selection cycle and the phase difference between the selection coefficient and population size. In particular, we find that the time-averaged SFS in slowly to moderately fast varying, on-average neutral environment has the same qualitative shape as the equilibrium SFS for positively selected mutant but differs quantitatively from it which can be captured by an effective population size.

## 1 Introduction

Despite several decades of intense research, the factors that determine the genetic variation in a population are not completely understood (Ellegren and Galtier, 2016; Moutinho *et al*., 2020). An important statistic for measuring the within-population genetic diversity is the site frequency spectrum (SFS) which gives the (unnormalized) distribution of allele frequency across polymorphic loci in the genome. It is known that in a neutral population of constant size, the equilibrium SFS decreases with the allele frequency of the derived allele whereas it is a U-shaped function when the mutant allele is under constant, positive selection (Wright, 1938; Kimura, 1968). However, these classical results can change due to complex evolutionary processes such as hitchhiking of neutral alleles with a beneficial mutant (Braverman *et al*., 1995; Fay and Wu, 2000), and in populations with high mutation rate (Desai and Plotkin, 2008; Charlesworth and Jain, 2014), fat-tailed offspring distribution (Birkner *et al*., 2013), seed bank (Koopmann *et al*., 2017), etc..

The above discussion assumes that the environment remains constant over evolutionary time scales. But in a changing environment, in general, there is no stationary state and the nonequilibrium SFS can differ significantly from the equilibrium one. For example, in an expanding neutral population, there is an excess of rare variants as compared to when the population size remains constant at its initial size, while population bottleneck leads to a deficit in the low frequency alleles due to the stronger effect of random genetic drift (Schraiber and Akey, 2015). The SFS is also related to other measures of genetic diversity such as mean heterozygosity and mean number of segregating sites which have been extensively studied when the population size is variable and selection is absent (Nei *et al*., 1975; Maruyama and Fuerst, 1985; Tajima, 1989), and in recent years, there has been some progress in understanding the effect of demography on SFS in a population under constant selection (Bustamante *et al*., 2001; Williamson *et al*., 2004, 2005; ŽivkoviĆ *et al*., 2015). However, the impact of changing selection on the nonequilibrium SFS is little studied (Huerta-Sanchez *et al*., 2008; Gossmann *et al*., 2014)) and to our knowledge, the *joint* effect of changing selection and demography has not been investigated so far.

Furthermore, most studies on the nonequilibrium SFS are concerned with historical demography such as population expansion in human population at the end of the last ice age (Williamson *et al*., 2005) or population bottleneck in non-African population of *D. melanogaster* around 6000 years ago (Baudry *et al*., 2004). However, variation in population size occurs over short time scales also for which different demographic models are needed. An example is the oscillatory change in the prey and predator population size over the course of a few generations (see, for e.g., Chapter 3 of Murray (2002)).

Motivated by the above discussion, here we consider a randomly mating, diploid population in which either population size and/or the selection coefficient change with time in a periodic fashion. We will explore the effect of selection strength, frequency of environmental variation and genetic dominance on the time-dependent SFS, and focus on how the time-averaged SFS deviates from the equilibrium SFS.

## 2 Model

We consider a randomly mating, diploid population of size *N* (*t*) at time *t* in which an individual’s genome is described by an infinite-sites model (Kimura, 1969) that makes the following key assumptions: first, a large number of sites are available for mutation which occurs with a small rate *μ* per site so that every mutation occurs (irreversibly) at a new site, and second, recombination is free so that at each site, allele frequency dynamics occur independently.

Furthermore, at each site, the homozygotes *AA* and *aa* have fitness 1 and 1+*s*(*t*), respectively, and the fitness of the heterozygote *aA* is 1+ *hs*(*t*) where the dominance coefficient 0 < *h* < 1. The frequency *x* of the mutant allele *a* increases or decreases at rate *r*_*b*_ or *r*_*d*_ respectively, which are given by (Devi and Jain, 2020; Kaushik and Jain, 2021)

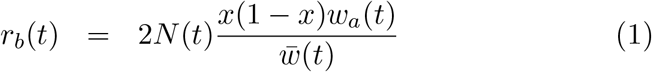

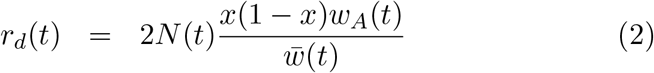

where *w*_*a*_(*t*) = (1 + *s*(*t*))*x* + (1 + *hs*(*t*))(1 − *x*) and *w*_*A*_(*t*) = (1 − *x*) + (1 + *hs*(*t*))*x* are the marginal fitness of allele *a* and *A*, respectively, and 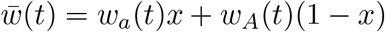 is the average fitness.

The population size and the selection coefficient are assumed to vary in time in a periodic fashion. We therefore write the selection coefficient as

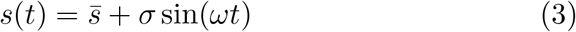

where the average 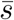 is arbitrary but the amplitude *σ* ≥ 0. The population size varies deterministically as

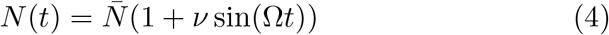

where 0 ≤ *ν* < 1 to ensure that the total population size is not subject to stochastic dynamics.

## 3 Site frequency spectrum: diffusion theory

To obtain an analytical understanding of the SFS and other measures of genetic diversity such as the mean heterozygosity, we work in the framework of the diffusion theory. It has been shown that in a large population, the average density *f* (*x, t*) of polymorphic sites with mutant allele frequency *x* at time *t* obeys the following partial differential equation: 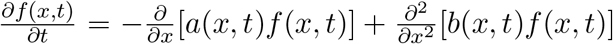 where *a*(*x, t*) and *b*(*x, t*) are, respectively, the mean and variance of the transition probability (Evans *et al*., 2007).

For the model detailed in Section 2, these coefficients have been derived in our earlier work (Kaushik and Jain, 2021), using which we find that

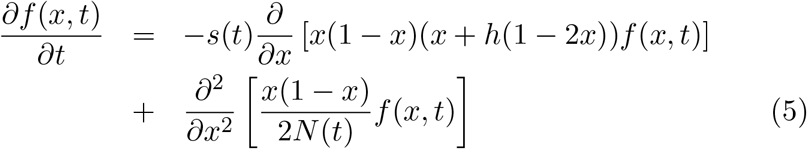

The above equation is subject to the initial condition *f* (*x*, 0), and boundary conditions, Lim_*x*→0_*xf* (*x, t*) = 2*N* (*t*)*μ* and *f* (1, *t*) finite (Evans *et al*., 2007). The first and second term on the RHS of (5) describe the effect of selection and random genetic drift, respectively, and the boundary condition at low allele frequency models the balance between the loss of polymorphism due to the absorption of the mutant allele by genetic drift alone and the gain by new mutations (for other modeling approaches, see ŽivkoviĆ *et al*. (2015); Gravel (2016)).

However, for both numerical and analytical purposes, it is easier to work with the density *g*(*x, t*) = *x*(1 − *x*)*f* (*x, t*) which obeys (Evans *et al*., 2007),

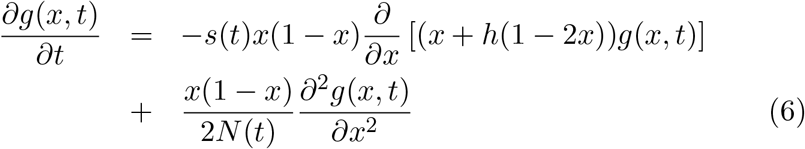

with initial condition *g*(*x*, 0) = *x*(1 − *x*)*f* (*x*, 0), and boundary conditions

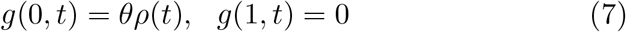

where 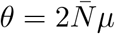 and 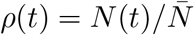. As discussed in Appendix A, one way to handle (6) is to carry out an eigenfunction expansion of *g*(*x, t*); unfortunately, the required eigenfunctions are not known in a closed form when selection is present (Kimura, 1957). However, as detailed in the following section, it is possible to make analytical progress even when the mutant allele is under selection in the limiting cases where the environment is changing either slowly or rapidly (Jung, 1993).

In the following, we will also study the unfolded sample frequency spectrum defined as the expected number of sites at which exactly 1 ≤ *i* ≤ *n* − 1 individuals in a population sample of size *n* ≪ *N* carry a mutation which is given by (Griffiths, 2003)

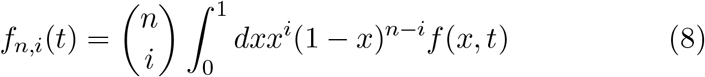

The sample SFS is also related to the mean heterozygosity,

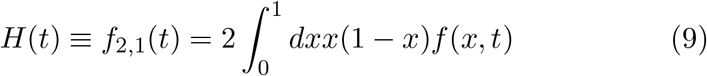

We will mainly focus on the time-averaged statistics obtained by averaging the time-dependent quantity *Q*(*t*) over the slower cycle, that is,

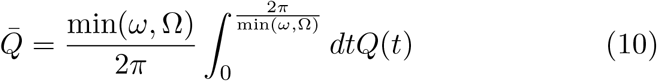

and compare the results with those in the constant environment to ascertain the effect of the changing environment.

## 4 Results

### 4.1 Static environment

Although the equilibrium SFS is known for a long time, it’s expression is not transparent and simple analytical approximations have not been obtained, and therefore, we first discuss these below. In a constant environment, polymorphic sites are lost due to absorption of the mutant allele and created by new mutations thus resulting in a steady state. Therefore, for 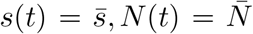, on setting the LHS of (5) equal to zero, the stationary state SFS is found to be (Wright, 1938, 1942; Kimura, 1968, 1969; Williamson *et al*., 2004)

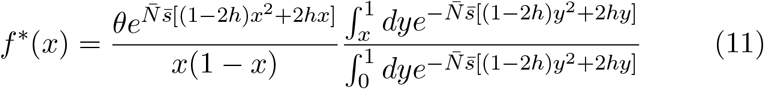

The above expression can also be derived by noting that as *θ* mutations occur per unit time per site in the population, *f* *(*x*)*dx*/*θ* is the mean sojourn time of the mutant allele between frequency *x* and *x* + *dx* before absorption, when the mutant allele is initially present in a single copy (Ewens, 2004).

#### 4.1.1 Weak selection

A power series expansion of (11) in scaled selection strength 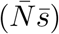 yields

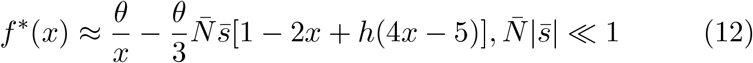

which, as for a neutral population, decreases monotonically with the allele frequency. This is because at most sites, single mutant will quickly get lost due to genetic drift, and therefore the average number of loci with very low mutant frequency would be high, and only a small number of loci will have intermediate to high allele frequency.

#### 4.1.2 Strong selection

As explained in Appendix B, when selection is strong, the integrals in (11) can be estimated using (B.2) for *h* ≠ 0, 1, and we obtain

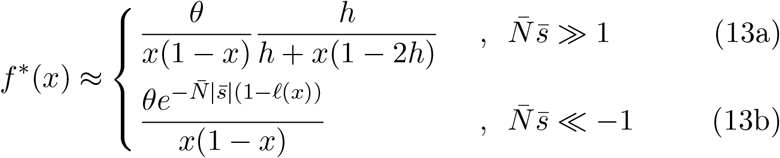

where *ℓ*(*x*) = (1 − *x*)[(1 − *x*) + 2*x*(1 − *h*)] (for completely dominant and recessive mutant allele, see Appendix B). Figures 1a and 1b show that when selection is strong, except for high allele frequencies, the exact equilibrium SFS given by (11) is in good agreement with the approximate expressions (13a) and (13b).

**Figure 1:**
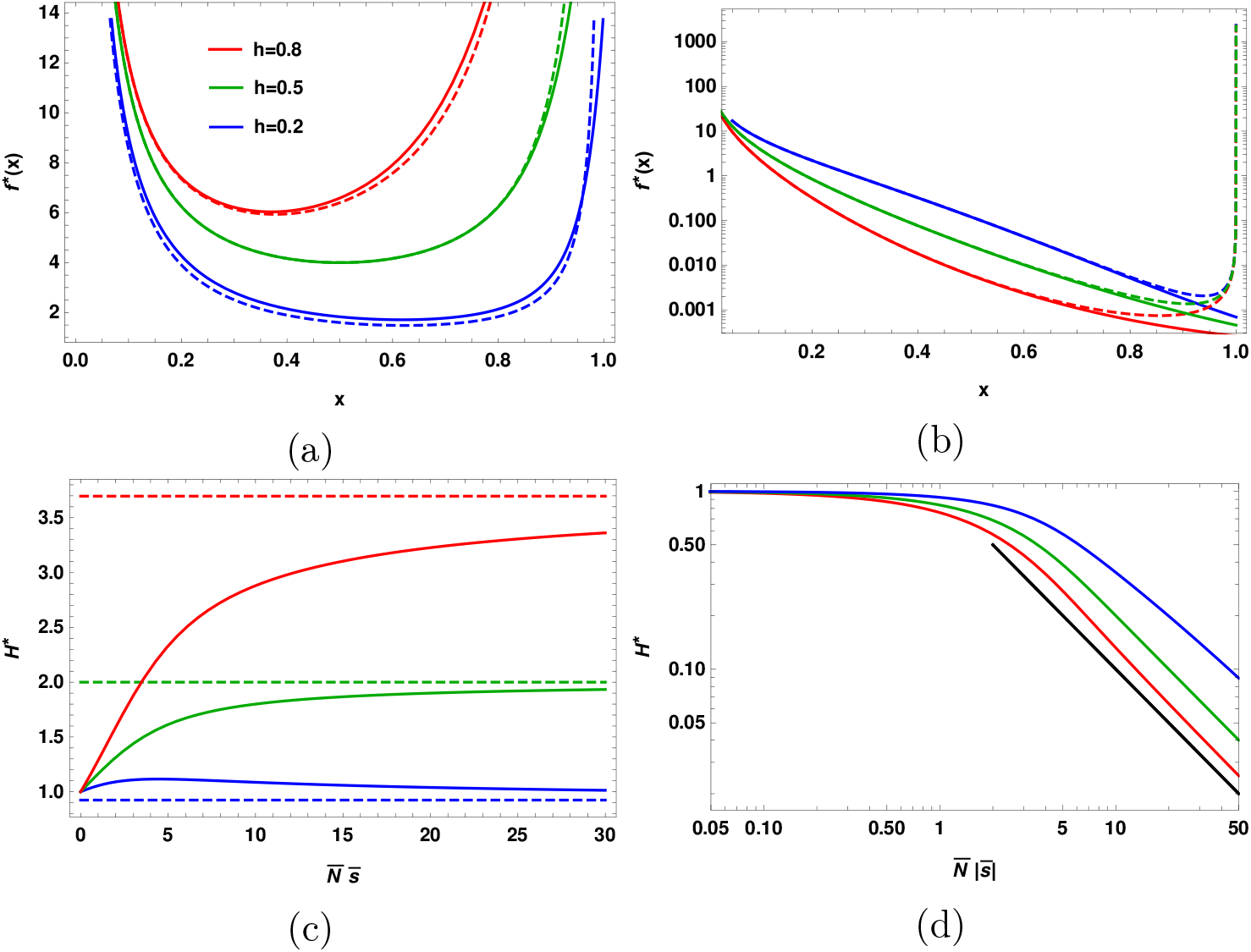
Static environment: The top panel shows the site frequency spectrum of a large population for selection strength (a) 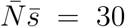, and (b) 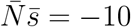. The solid lines show the exact expression (11), while the dashed lines show the approximate expression (13a) and (13b) for beneficial and deleterious mutations, respectively. The bottom panel shows the mean heterozygosity calculated numerically using (11) for (c) positively selected mutant which approaches the asymptotic value (15b) depicted by dashed lines, and (d) negatively selected mutant that decreases towards zero as 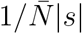 (black curve) for all *h* in accordance with (15c). In all the plots, the scaled mutation rate *θ* = 1.

Equations (13a) and (13b) show that for 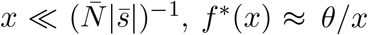 so that at low allele frequencies, the SFS for strong selection behaves the same way as that for a neutral population, and the effect of selection becomes apparent only at intermediate and high allele frequencies. For negative selection, the SFS decreases monotonically towards zero with the mutant allele frequency for the same reason as for the neutral population. But for positive selection, it has the characteristic U-shape reflecting the fact that the mean sojourn time is long at high allele frequencies for beneficial mutations (Ewens, 2004).

From (13a) and (13b), we also find that for a given allele frequency, the SFS increases with the dominance coefficient for beneficial mutations and decreases for deleterious ones; this is due to the longer sojourn time to fixation for dominant, advantageous mutation and the fact that the deleterious recessives are more likely than the dominant alleles to reach finite frequencies (Williamson *et al*., 2004).

#### 4.1.3 Sample SFS

Using the approximate results (12), (13a) and (13b) for a large population in (8), we find the sample SFS to be (see Appendix B for details)

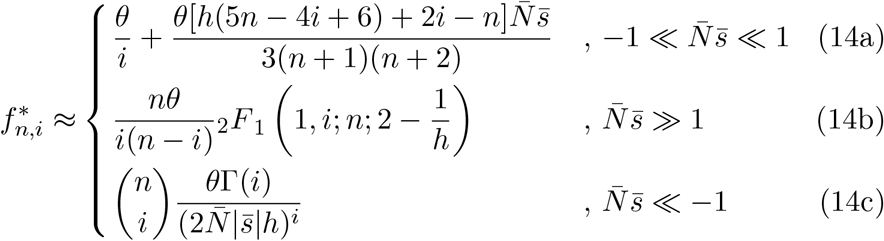

where Γ(*i*) is the gamma function and _2_*F*_1_(*a, b*; *c*; *z*) is the Gauss hypergeometric function (Abramowitz and Stegun, 1964). For *i* ≪ *n* − 1, the contribution to the integral on the RHS of (8) comes from small *x*, while for large *i*, variants at high frequency matter; as a result, the sample SFS behaves qualitatively in the same manner as the SFS for a large population discussed above.

The neutral mean heterozygosity in a constant size population is given by *θ*. From (14a)-(14c), we find that when the mutant allele is under selection, the mean heterozygosity can be approximated by

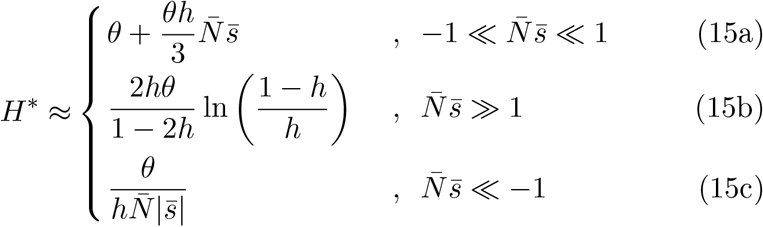

As positive (negative) selection decreases (increases) the chance of loss of the new mutant allele, one expects the heterozygosity to be enhanced (reduced) with increasing magnitude of selection for beneficial (deleterious) mutants; indeed, (15a) and (15c) are consistent with this expectation. But, for strong positive selection, as Figure 1c shows, the mean heterozygosity does not necessarily increase with selection. In fact, on equating the RHS of (15b) to *θ*, we find that the mean heterozygosity saturates to a value smaller than that for the neutral allele for 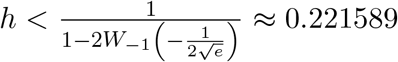 where *W*_−1_(*z*) is the lower branch of the Lambert W function (Corless *et al*., 1996). We also note that with increasing magnitude of selection, the mean heterozygosity approaches the asymptotic value, viz., (15b) for beneficial mutant and zero for deleterious one, algebraically slowly (see (B.6a) and (B.6b), and Fig. 1d).

For positive selection, on comparing (15b) with the mean heterozygosity 2*θ* for the co-dominant mutant, we find that *H*\* for a dominant (recessive) allele is larger (smaller) than that for the co-dominant allele; the mean heterozygosity increases with the dominance level since the mutant’s chance of escaping stochastic loss is enhanced with increasing *h* (Haldane, 1927), and hence it can contribute to the population diversity before eventually fixing. Similarly, one can argue that for negative selection, *H*\* will decrease with increasing *h* (see Figs. 1c and 1d).

### 4.2 Slowly changing environment

We now turn to the properties of SFS in slowly changing environments. If the frequency with which the selective environment and population size change is much smaller than the inverse average population size and average selection coefficient (that is, 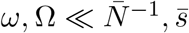), one can obtain the time-dependent SFS within an *adiabatic approximation* (Jung, 1993). For this purpose, we first rewrite (5) as

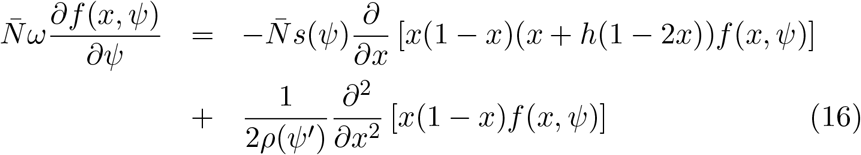

where *ψ* = *ωt, ψ*′ = Ω*t* and the ratio 0 < Ω/*ω* < ∞. For arbitrary scaled selection, on expanding *f* (*x, ψ*) as a power series in 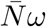 and substituting it in the above equation, we find that to leading order in 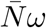, the LHS of (16) is zero. It is easy to verify that this result also holds when either the population size or the selection coefficient varies with time. Thus in slowly changing environment, the mean density *f* (*x, t*) of sites with allele frequency 0 < *x* < 1 obeys the steady state equation at *instantaneous N* (*t*) and *s*(*t*), and one can therefore obtain *f* (*x, t*) by simply letting 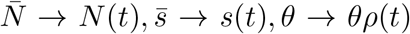 in (11) and arrive at

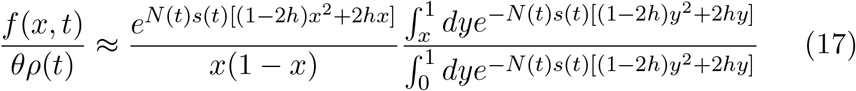

Figure 2a shows that (17) is in good agreement with the results obtained by numerically integrating (5) for small cycling frequencies. We find that for strong positive selection, *f*(*x, t*) is nonzero and increases (decreases) with increasing (decreasing) selection strength but it is close to zero when the selection strength is small or negative; these behavior may be understood by appealing to the results (13a) and (13b) in static environments. It is evident from (17) that when either *N* or *s* is time-dependent, *f*(*x, t*) is a periodic function in time with the period of the varying parameter; if both *N* and *s* vary such that *ω* = (*n*_1_/*n*_2_)Ω with *n*_1_, *n*_2_ being integers and *n*_1_ < *n*_2_ (say), the period of *f*(*x, t*) is given by 2*πn*_1_/Ω.

**Figure 2:**
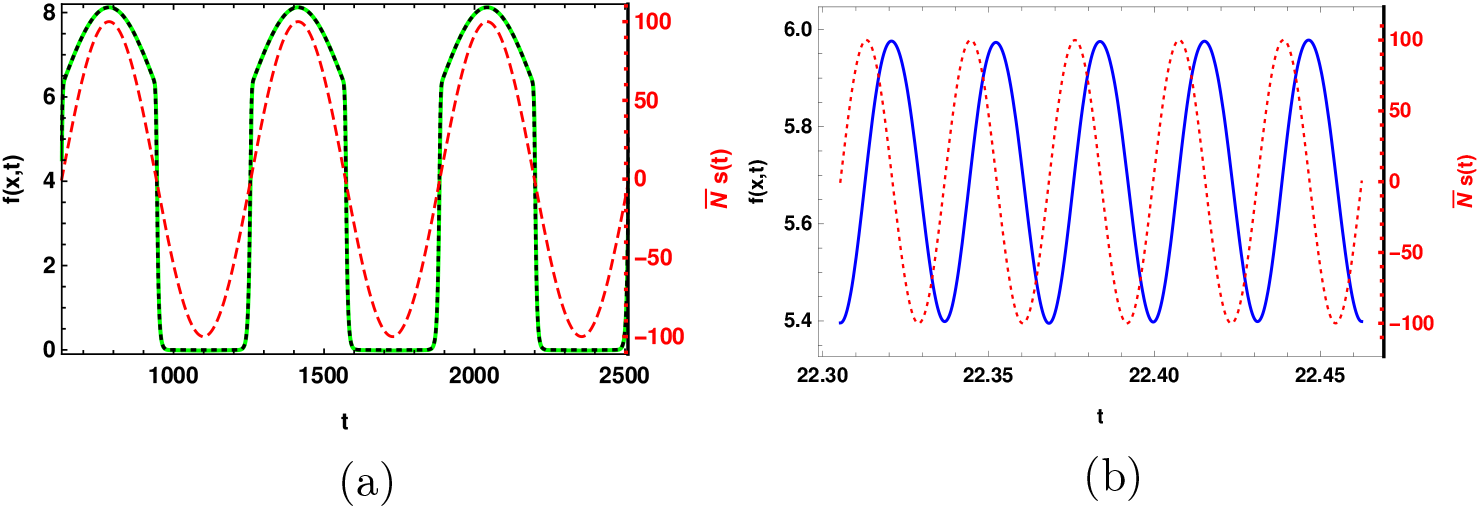
Changing environment and on-average neutral selection: The time-dependent site frequency spectrum (solid line) in a large population when the population size and selection coefficient (dotted line) change periodically in time with equal cycling frequency obtained by numerically solving (5) for (a) 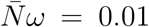 and (b) 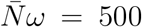. The result (17) obtained within adiabatic approximation (dashed line) is also shown for comparison in the slowly changing environment. The other parameters are *x* = 0.2, *ν* = 0.3, *h* = 1/2, *θ* = 1 and 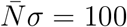.

#### 4.2.1 No selection

In the absence of selection, (17) gives *f*(*x, t*) = *θρ*(*t*)/*x* from which it immediately follows that the time-averaged SFS, 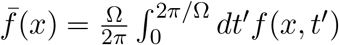 is simply *θ*/*x* and the time-averaged sample SFS is equal to *θ*/*i*. In Appendix C, these results are obtained using an eigenfunction expansion of *f*(*x, t*) which furthermore shows that the correction to the adiabatic approximation is of order 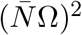, see (C.17). Figure 3 shows the sample SFS for various Ω’s, and we find that it is well approximated by *θ*/*i* when the population size changes slowly.

**Figure 3:**
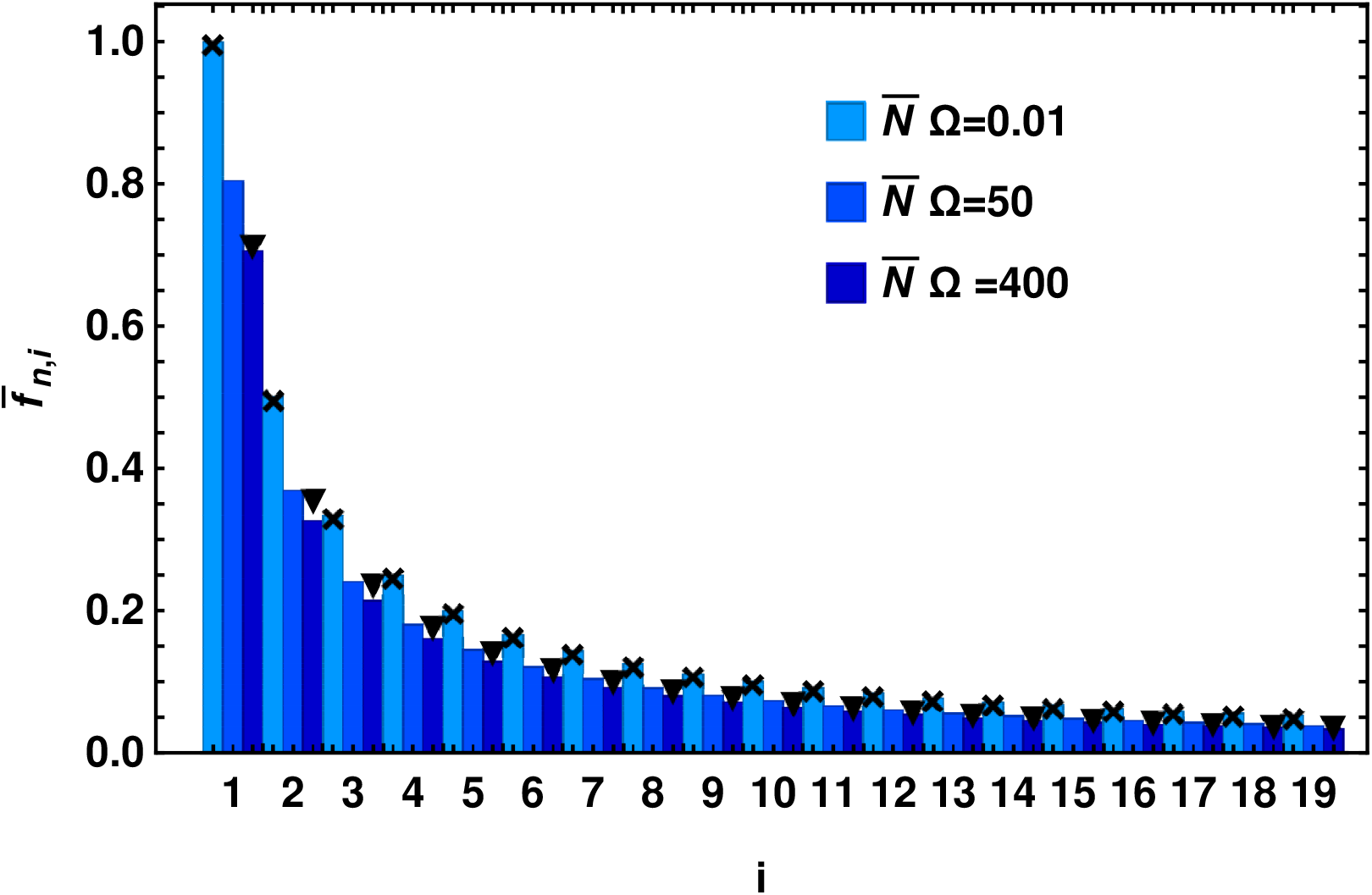
Changing population size and no selection: The time-averaged sample site frequency spectrum when population size changes with time for sample size *n* = 20 and dominance coefficient *h* = 1/2. The bars are obtained by numerically solving (5) and (8) for different cycling frequencies while the points show (C.17) and (C.24) for small and large cycling frequencies, respectively.

#### 4.2.2 Weak selection

When both population size and selection vary with the same frequency, using (10) and (12), we find that the time-averaged SFS, 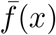 differs from *f*\*(*x*) by an amount

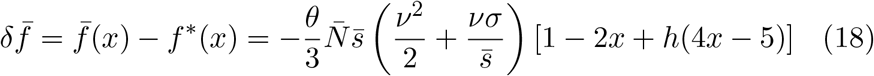

Similarly, the change in the time-averaged mean heterozygosity 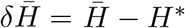 due to slowly changing environment is given by

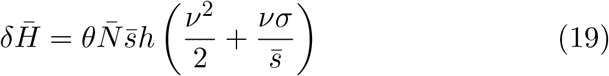

The factor in the bracket on the RHS of the above equation captures the joint effect of the change in the selection and population size, and shows that (to leading order in 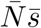), changing selection but constant population size results in the same heterozygosity as in the static environment and therefore the mean heterozygosity is affected mainly due to demography; however, as expected, the change 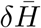 is small for weak selection, as shown in Fig. 4a and Fig. 5a.

**Figure 4:**
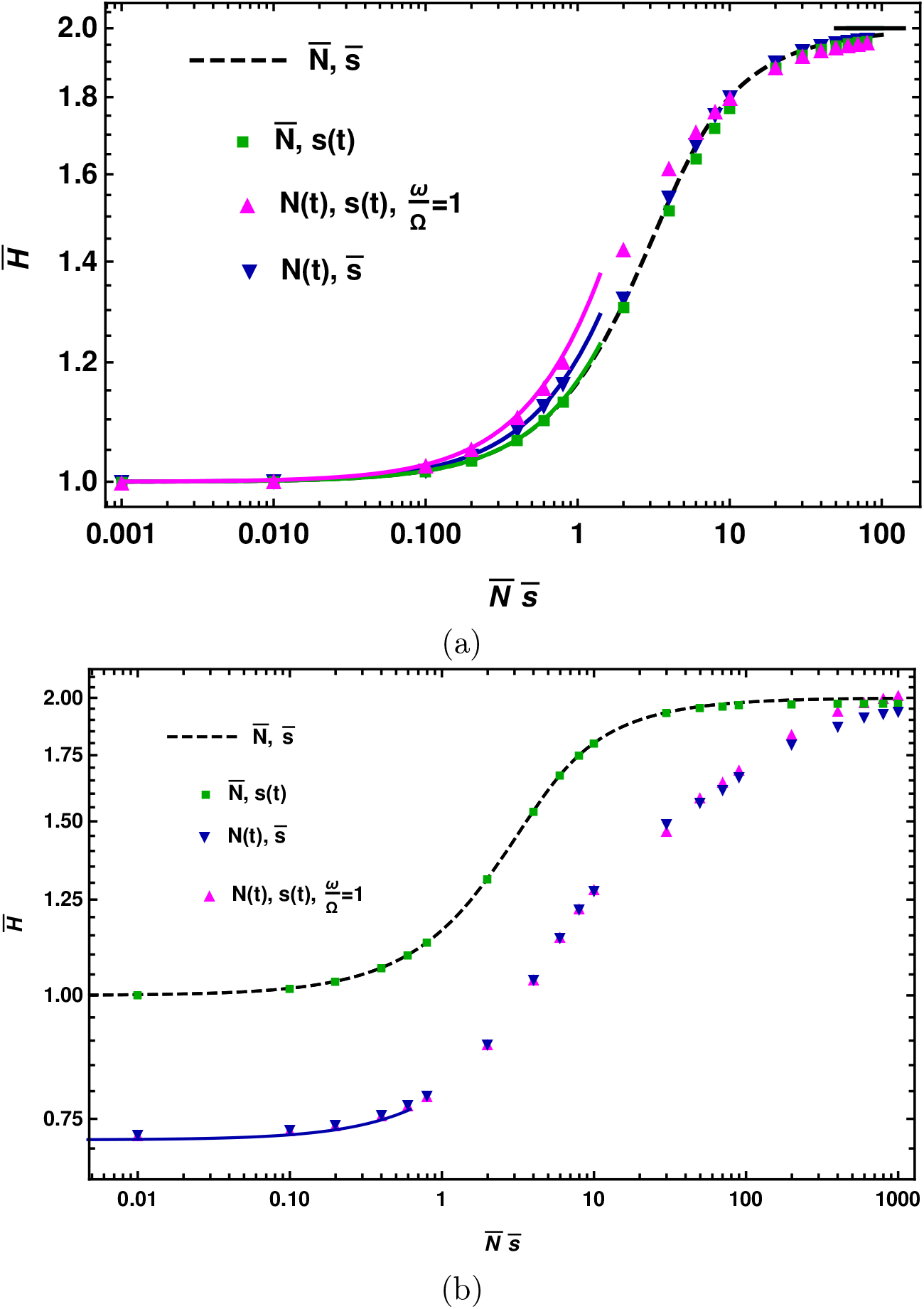
Changing environment and positive selection at all times: The time-averaged heterozygosity, 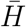 as a function of selection strength when either the selection coefficient (indicated in the legend by 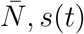) or population size 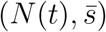 or both (*N*(*t*), *s*(*t*)) vary with time. The top panel shows the results in slowly changing environments 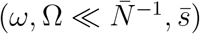 for 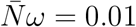, and the bottom panel in rapidly changing environments 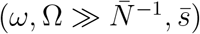 for 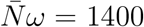. The points are obtained by numerically solving (5) and (9) for different scenarios as indicated in the legend for 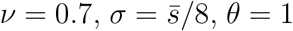, and *h* = 1/2. The broken line shows the results in the static environment that are obtained using (11). The solid lines show the analytical approximation (20) and (24) for weak and strong selection, respectively, in slowly changing environment, and (34) for weak selection in rapidly changing environment.

**Figure 5:**
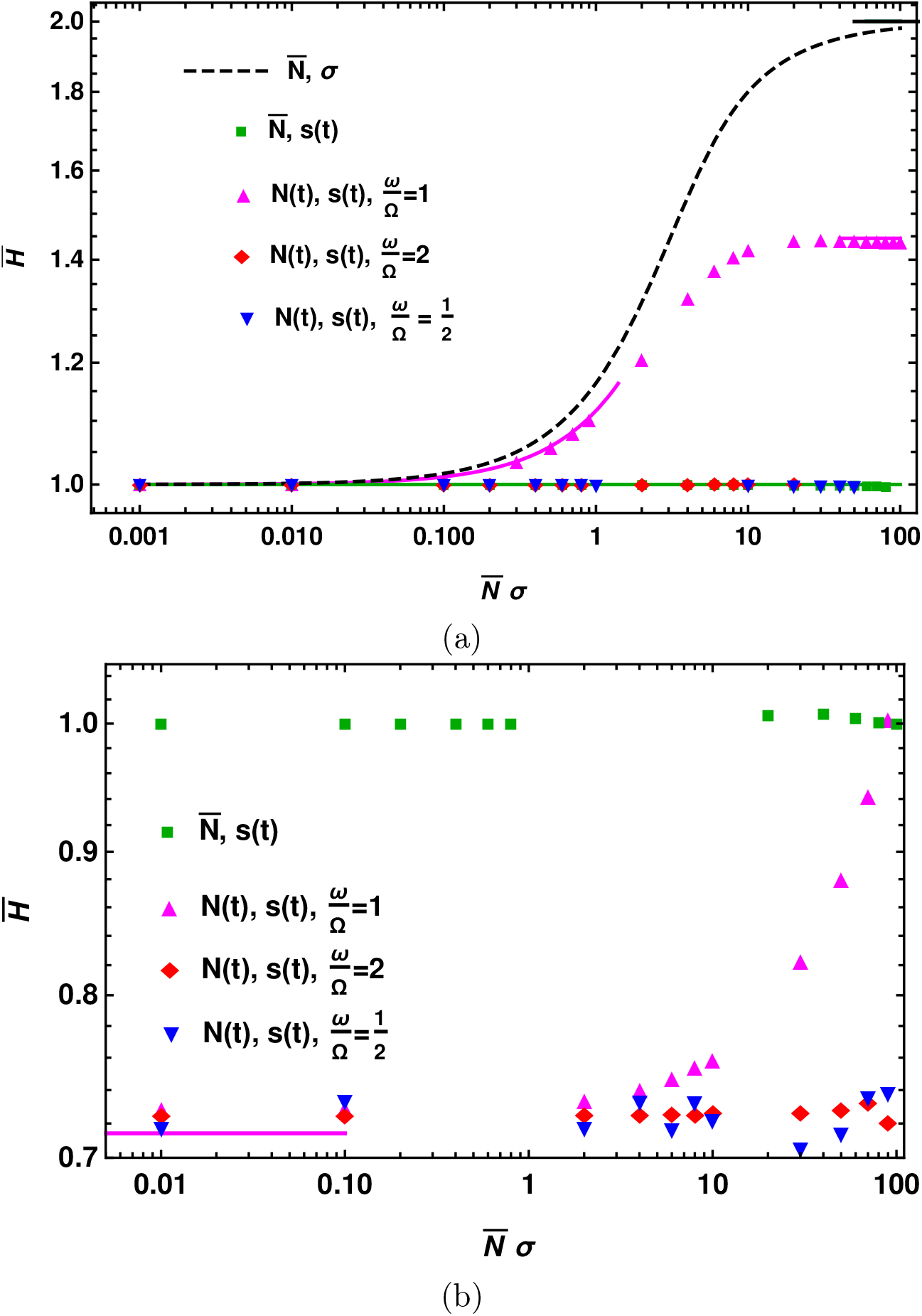
Changing environment and on-average neutral selection: The time-averaged mean heterozygosity, 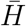 as a function of selection strength when either the selection coefficient (indicated in the legend by 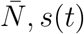) or population size 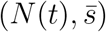 or both (*N*(*t*), *s*(*t*)) vary with time. The top panel shows the results in slowly changing environments 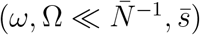 for 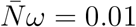, and the bottom panel in rapidly changing environments 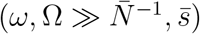 for 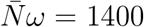. The points are obtained by numerically solving (5) and (9) for different scenarios as indicated in the legend for *ν* = 0.7, *θ* = 1, *h* = 1/2 and no phase difference between selection and population size variation. The broken line shows the results in the static environment that are obtained using (11). The solid lines show the analytical approximation (20) for weak selection, and (27) and (28) for strong selection in slowly changing environment.

The correction to the adiabatic approximation is obtained in Appendix D when selection is weak and constant but population size varies, and we find that the time-averaged mean heterozygosity is given by (see (D.8) and (D.11)),

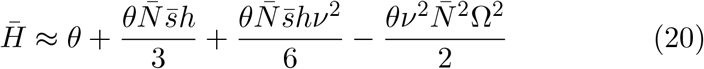

The first three terms on the RHS of the above equation match with the result from adiabatic theory, and the last term can be ignored for 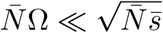.

#### 4.2.3 Strong, on-average nonzero selection

On replacing the selection coefficient and population size by the corresponding time-dependent quantities in (13a) and (13b), we obtain the time-dependent SFS for on-average non-neutral mutants under strong selection to be

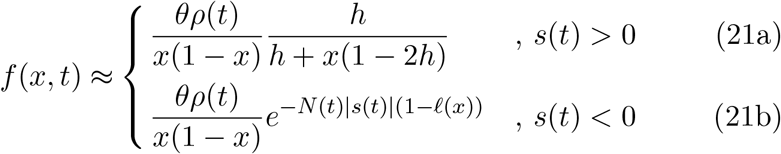

If both selection and population size change with the same frequency and selection is strong enough so that the exponentially small contribution in selection strength from the negative cycle of selection can be ignored, then the time-averaged SFS is given by

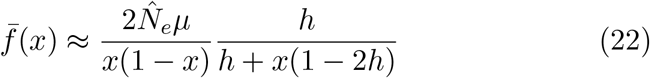

where

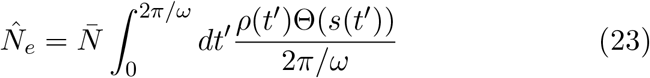

and Θ(*x*) is the Heaviside step function. The effective population size 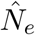 defined above states that the contribution to 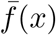 comes from the average population size during the time selection is positive.

This shows that when both selection and population size are changing but selection remains positive throughout the cycle, 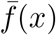 is well approximated by the static result given by (13a). Figure 4a shows the time-averaged heterozygosity when the selection is positive at all times which, in the light of the above discussion, is given by

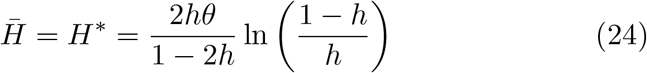

However, for 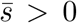, if *s*(*t*) is negative for a part of the cycle, (22) shows that 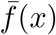 is smaller than *f*\*(*x*). Thus, in a slowly changing environment, for strong selection, the SFS is reduced if the mutant becomes deleterious during a part of the cycle. Similarly, for 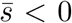, the SFS is enhanced due to nonzero contribution from the positive part of the selection cycle.

#### 4.2.4 Strong, on-average zero selection

One can obtain explicit expressions for the time-averaged SFS when the mutant spends equal time in both positive and negative part of the selection cycle (that is, 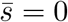). For 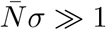, from (22), we get

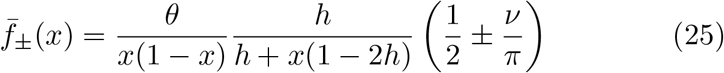

for *ρ*_±_(*t*) = 1 ± *ν* sin(*ωt*). Thus we find that unlike in the static, neutral environment where the SFS decays with allele frequency, the time-averaged SFS in an on-average neutral environment is a non-monotonic function of the allele frequency as, up to a scale factor, it is given by the equilibrium SFS for positively selected mutant (see (13a)).

The factor in the parentheses on the RHS of (25) shows that when the population size varies in-phase with selection, as the population size is larger than the average 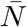 during the positive cycle and stronger positive selection leads to larger heterozygosity, demography ameliorates the effect of deleterious mutations. On the other hand, when selection and population size change out-of-phase, the mean heterozygosity is smaller than in the situation where only selective environment is varied. Quantitatively, from (25), we find that the relative mean heterozygosity 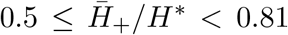 and 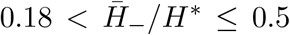 for 0 ≤ *ν* < 1.

When the population size is constant (*ν* = 0), selection is strong 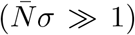 and *h* = 1/2, from (25), we find that the time-averaged mean heterozygosity is the same as in the constant, neutral environment. In fact, as shown in Appendix E, this result holds exactly for any cycling frequency and selection strength provided the population size remains constant:

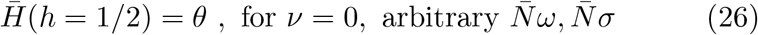

However, for *h* ≠ 1/2, from (15b) and (25), we find that 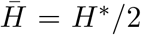 which is smaller than *θ* for *h* < 1/2 and larger for *h* > 1/2.

We now consider the general situation where the population size and selection coefficient vary with different cycling frequencies and have a phase difference. For simplicity, we assume that the ratio of the two frequencies is an integer and consider the average over the smaller frequency (that is, larger time period) as defined in (10). If selection changes slower than the population size (Ω = *Kω, K* = 1, 2, …), and *s*(*t*) = *σ* sin(*ωt*), *ρ*(*t*) = 1 + *ν* sin(Ω*t* + Φ), 0 ≤ Φ < 2*π*, averaging over the selection cycle in (22), we obtain

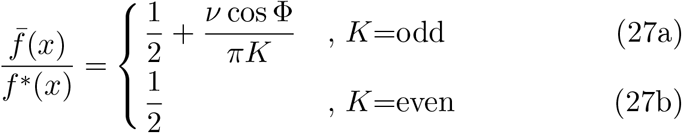

To understand the *K*-dependence, we consider the case when Φ = 0. When an even number of cycles of population size occur during one cycle of selection, *K*/2 complete cycles of *N* (*t*) lie in the beneficial part of the selection cycle. But as the average population size in each of the *K*/2 cycles is 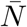, the final result (27b) is found to be independent of *K*. In contrast, for odd *K*, the first one half of the 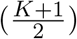 th cycle of *N* (*t*) contributes more than 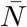 but it’s duration shrinks with increasing *K* as captured by the second term on the RHS of (27a). Similarly, when selection changes faster than the population size (*ω* = *κ*Ω, *κ* = 1, 2, …), and *s*(*t*) = *σ* sin(*ωt*), *ρ*(*t*) = 1 + *ν* sin(Ω*t* + *ϕ*), 0 ≤ *ϕ* < 2*π*, we obtain

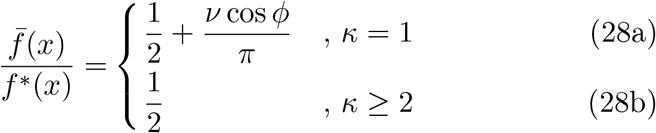

The above discussion thus demonstrates that unequal cycling frequencies and phase difference can affect the time-averaged SFS. Figure 5a summarizes the results obtained above, and shows that for strong selection, the time-averaged mean heterozygosity in slowly changing environment is smaller than that in the constant environment with population size 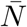 and constant, positive selection coefficient *σ* but it can be equal to that in the constant neutral environment. (Note that 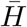 can be smaller than *θ* if population size and selection coefficient variation have a phase difference.)

From (14b) and (14c), for in-phase variation in population size and selection coefficient with equal cycling frequency, we find the time-averaged sample SFS to be

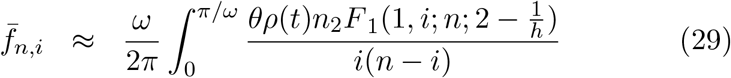

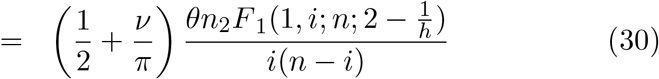

Figure 6a shows that in slowly changing environment, the sample SFS obtained by numerically integrating (5) and the above analytical approximation are in good agreement except at *i* = 1, *n* − 1. Thus the sample SFS in on-average neutral environment is not a monotonically decreasing function unlike for equilibrium neutral SFS. In Fig. 6a, the nonequilibrium SFS is also compared with the equilibrium SFS for mutant under strong, positive selection with selection strength 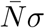, and we find that the sample SFS in changing environment is consistently smaller than the equilibrium SFS.

**Figure 6:**
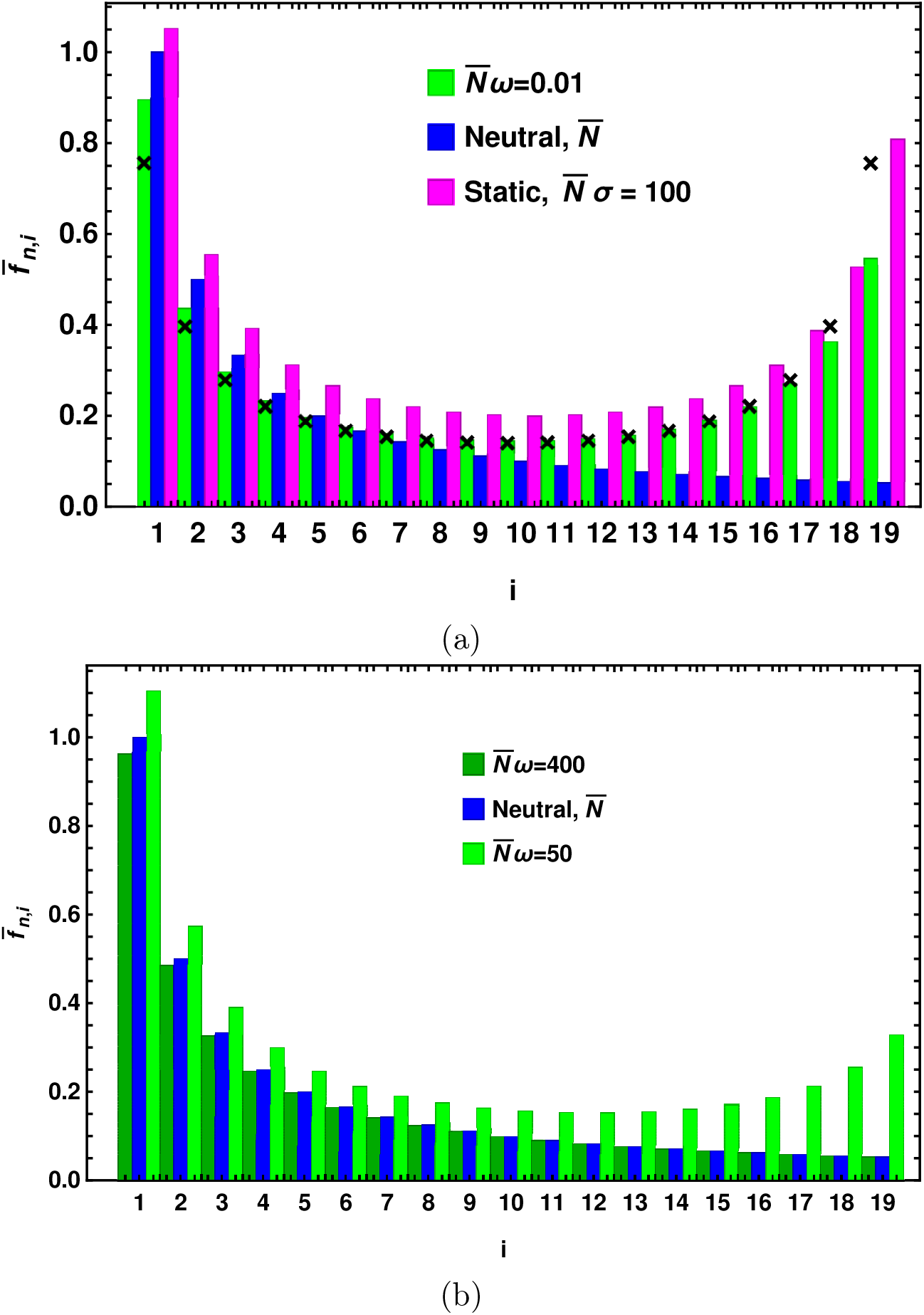
Changing environment and on-average neutral selection: The time-averaged sample site frequency spectrum represented by bar chart and obtained by numerically solving (5) and (8). The top panel shows the results in slowly changing environment for 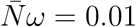 that are compared with equilibrium SFS for neutral and positively selected mutant in constant environment. The bottom panel shows the sample SFS in rapidly changing environments for two cycling frequencies which is compared with the equilibrium neutral SFS. The points in the top panel are obtained using (30). The other parameters are 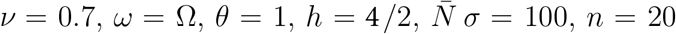 and no phase difference between selection and population size variation.

### 4.3 Rapidly changing environment

We now consider the parameter regime where the rate of change of population size and selection is much larger than the inverse population size and average selection coefficient 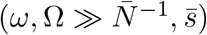. Figure 2b shows the periodic variation of the time-dependent SFS with changing environment for large frequencies. We note that, unlike in slowly changing environment, there is a phase difference between the SFS and environment: the SFS increases while the selection is positive and decreases in the deleterious part of the cycle.

In rapidly changing environments, a general expectation is that the population is sensitive only to the time-averaged environment (SjÖdin *et al*., 2005). If the population size remains constant in time but selection varies rapidly, the boundary condition (7) for *g*(0, *t*) is time-independent, and one can use a standard argument to find 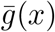 (Reimann *et al*., 1996). For this purpose, we first rewrite (6) as

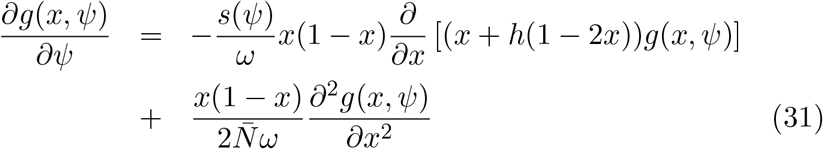

where *ψ* = *ωt*, and expand *g*(*x, ψ*) as a power series in 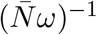, that is, write 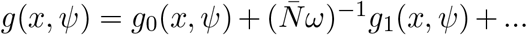, and substitute this expansion in the above equation. Keeping terms to leading order in the expansion parameter, we obtain 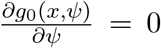, which implies that *g*_0_ is a function of *x* alone and subject to boundary conditions *g*_0_(0) = *θ, g*_0_(1) = 0. As a result, the next order term *g*_1_ obeys the following partial differential equation:

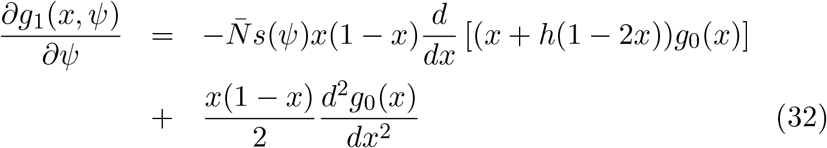

Integrating both sides of the above equation over the period 2*π* and using the periodicity property of *g*(*x, ψ*), we find that the LHS of the above equation is zero, and *f*_0_(*x*) = *g*_0_(*x*)/(*x*(1 − *x*)) is given by (11).

Therefore, when the population size is constant, the time-averaged SFS in a rapidly varying environment is simply given by the equilibrium SFS in the time-averaged environment. For this reason, the time-averaged mean heterozygosity shown in Fig. 4b and Fig. 5b when only selection changes matches with the equilibrium mean heterozygosity in the time-averaged environment for any (nonnegative) selection strength. But when demographic changes also occur, as the boundary condition *g*(0, *t*) is time-dependent, it is not clear how to generalize the above argument. As we will discuss below, the time-averaged selective environment is the primary determinant of the behavior of the time-averaged SFS: if the selective environment is neutral or weak on-average, the demography has a strong impact, but if selection is on-average strongly positive, changing population size does not influence the genetic diversity.

#### 4.3.1 No selection

Previous numerical studies (Nei *et al*., 1975; Maruyama and Fuerst, 1985) with demography and no selection 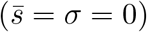 suggest that for large 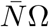, the time-averaged mean heterozygosity is equal to 2*N*_*e*_*μ* where *N*_*e*_ is the effective population size given by the harmonic mean over the period 2*π*/Ω,

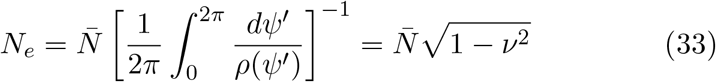

which decreases with increasing amplitude of the population size variation. In Appendix C, we show that at large cycling frequencies, indeed the effect of demography is to replace 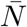 by *N*_*e*_ in the static environment results. Thus the time-averaged sample SFS is given by 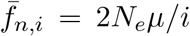 (see (C.24)) from which it immediately follows that the time-averaged mean heterozygosity, 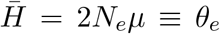. Figure 3 shows that the sample SFS is well approximated by *θ*_*e*_/*i* when the population size changes rapidly.

#### 4.3.2 Weak selection

In Appendix D, we develop a perturbation theory to understand the SFS in rapidly changing environments. In particular, for weak constant selection 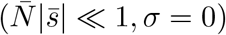, from (D.8) and (D.12), we find that the time-averaged mean heterozygosity is given by

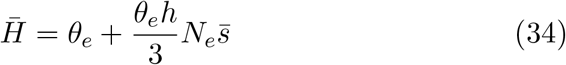

which is simply the equilibrium result (15a) for weakly selected mutant with population size *N*_*e*_ and matches with the numerical results shown in Fig. 4b. We have verified analytically that for weakly selected, on-average neutral mutant also, the effect of changing selection and demography is to replace 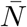 by *N*_*e*_ in the constant neutral environment results, and yield 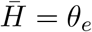 (refer to Fig. 5b).

Thus the above discussion suggests that when the environment changes rapidly and time-averaged selection is neutral or weak (while the population size may or may not vary), the time-averaged sample SFS is given by the expression (14a) in the constant environment with selection coefficient 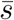 and population size *N*_*e*_,

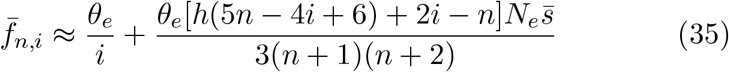

#### 4.3.3 Strong, on-average positive selection

For strong and positive selection 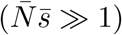, from (6), we have

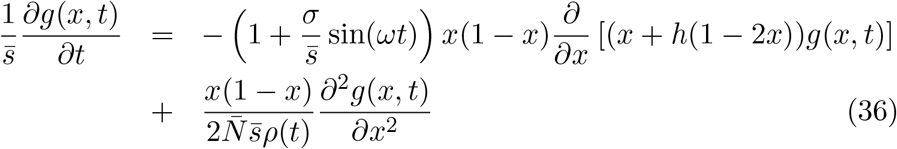

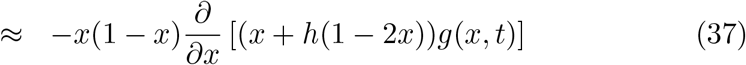

where the last equation is obtained for large 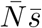 and 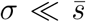. Then averaging both sides of (37) over the slower cycle (as defined in (10) and for integer ratio of the two cycling frequencies), and using the boundary condition 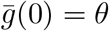, we obtain

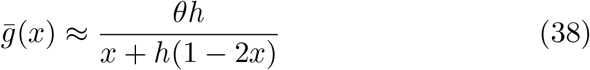

which gives the time-averaged SFS to be the same as for the positively selected mutant in constant environment with population size 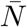 (see (13a)). Note that the above argument is valid for arbitrary scaled cycling frequency. Furthermore, due to (14b), the time-averaged sample SFS is given by

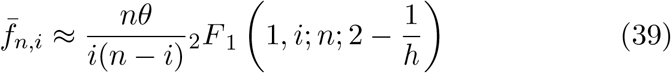

The above discussion shows that for a co-dominant mutant under strong selection, the time-averaged mean heterozygosity is equal to 2*θ* in both slowly and rapidly changing environments in agreement with the data shown in Figs. 4a and 4b. Thus, when both selection and population size vary rapidly, while the results for weak selection are given by the equilibrium ones on replacing 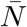 by *N*_*e*_, the equilibrium SFS for population size 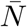 remains valid when selection is strong. This is because in the above deterministic analysis for large 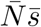, the genetic drift term on the RHS of (36) whose coefficient depends on the inverse population size is neglected, and the population size enters the expression for SFS through the scaled mutation rate only.

#### 4.3.4 Strong, on-average zero selection

When the selective environment is on-average neutral, the time-averaged mean heterozygosity displayed in Fig. 5b for large cycling frequencies is equal to *θ* for constant population size and changing selection as discussed at the beginning of Section 4.3. When both population size and selection vary with equal frequencies, 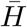 again approaches *θ*; however, for unequal frequencies, the results obtained by numerically integrating (5) suggest that 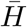 is independent of the selection strength and given by *θ*_*e*_. We do not have an analytical understanding of these results; see, however, Appendix F. Thus, barring the case when the cycling frequencies are equal, the time-averaged heterozygosity is given by that in the time-averaged selective environment for both weak and strong selection strength and hence the results obtained in Sec. 4.3.1 apply.

## 5 Discussion

In this article, we are interested in understanding how changing environment, be it due to demography and/or time-dependent selection, impacts genetic diversity within a population. The importance of demography in understanding various measures of genetic diversity has long been appreciated (Nei *et al*., 1975; Maruyama and Fuerst, 1985; Tajima, 1989; Williamson *et al*., 2005; ŽivkoviĆ *et al*., 2015), but the effect of fluctuating selection on genetic variation has been relatively less studied (Huerta-Sanchez *et al*., 2008; Gossmann *et al*., 2014). Here we have considered the effect of periodic changes in selection coefficient and population size on the site frequency spectrum and mean heterozygosity.

When selection is weak, as expected, the time-averaged mean heterozygosity is essentially the same as for a neutral population with changing population size (see Figs. 4 and 5). We find that the effect of demography is mainly seen when the population size changes rapidly where the time-averaged statistical quantities behave the same way as in the constant environment but with reduced population size, *N*_*e*_ given by the harmonic mean of the changing population size over a period (see (33)). Although the effective population size has been derived when the population size has random fluctuations, and verified numerically for other models of demographic change (Nei *et al*., 1975; Maruyama and Fuerst, 1985; Tajima, 1989), here we have analytically derived the effective population size for periodically changing population size in Appendices C and D. Our explicit expressions for the time-averaged SFS are given by (C.17) and (C.24) for slowly and rapidly changing environments, respectively, and shown graphically in Fig. 3 when selection is absent.

For strong selection, if time-averaged selection is positive, a deterministic argument, given in Sec. 4.3.3, shows that the time-averaged SFS is the same as for positively selected mutant in constant environment for any cycling frequency. But the results depend on the cycling frequency when the selection is zero on-average. For large cycling frequencies, the time-averaged mean heterozygosity is found, in general, to behave as in constant neutral environment with effective population size *N*_*e*_ discussed above; however, in slowly changing environment, the time-averaged SFS is the same as in the static environment with an effective population size 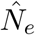 given by the average population size over the time the selection is positive, see (23). For neutral populations, the census population size in changing environment can be replaced by an appropriately defined effective population size when it’s mutation rate relative to the frequency of change in the population size is small (Nei *et al*., 1975) and stochastic changes in the population size are independent random variables (SjÖdin *et al*., 2005; Iizuka, 2010). Here, in addition to the demographic *N*_*e*_ in fast changing environment discussed above, we find that the census population size in slowly changing, selective environment can also be interpreted as an effective population size, 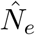 (refer to (23)).

Furthermore, in slowly changing, on-average neutral environment, we find that the time-averaged SFS is a U-shaped function as one would expect for beneficial mutants in constant environment. However, as shown in Fig. 6, relative to the SFS in constant environment, the time-averaged SFS is reduced by a factor equal to the fraction of the time the selection is negative in a seasonal cycle. Figure 6 also shows that there are more high frequency variants in the changing environment as compared to in the constant, neutral environment; similar behavior has also been seen when there is fluctuating selection (but no demography) (Huerta-Sanchez *et al*., 2008; Gossmann *et al*., 2014). Taken together, these studies suggest that varying selection, even if zero on-average, can produce a SFS similar to that as in constant, positively selective environment. Fig. 7 further shows that the time-averaged heterozygosity follows the same qualitative trend with dominance coefficient as for positively selected mutant in constant environment (see Fig. 1c).

**Figure 7:**
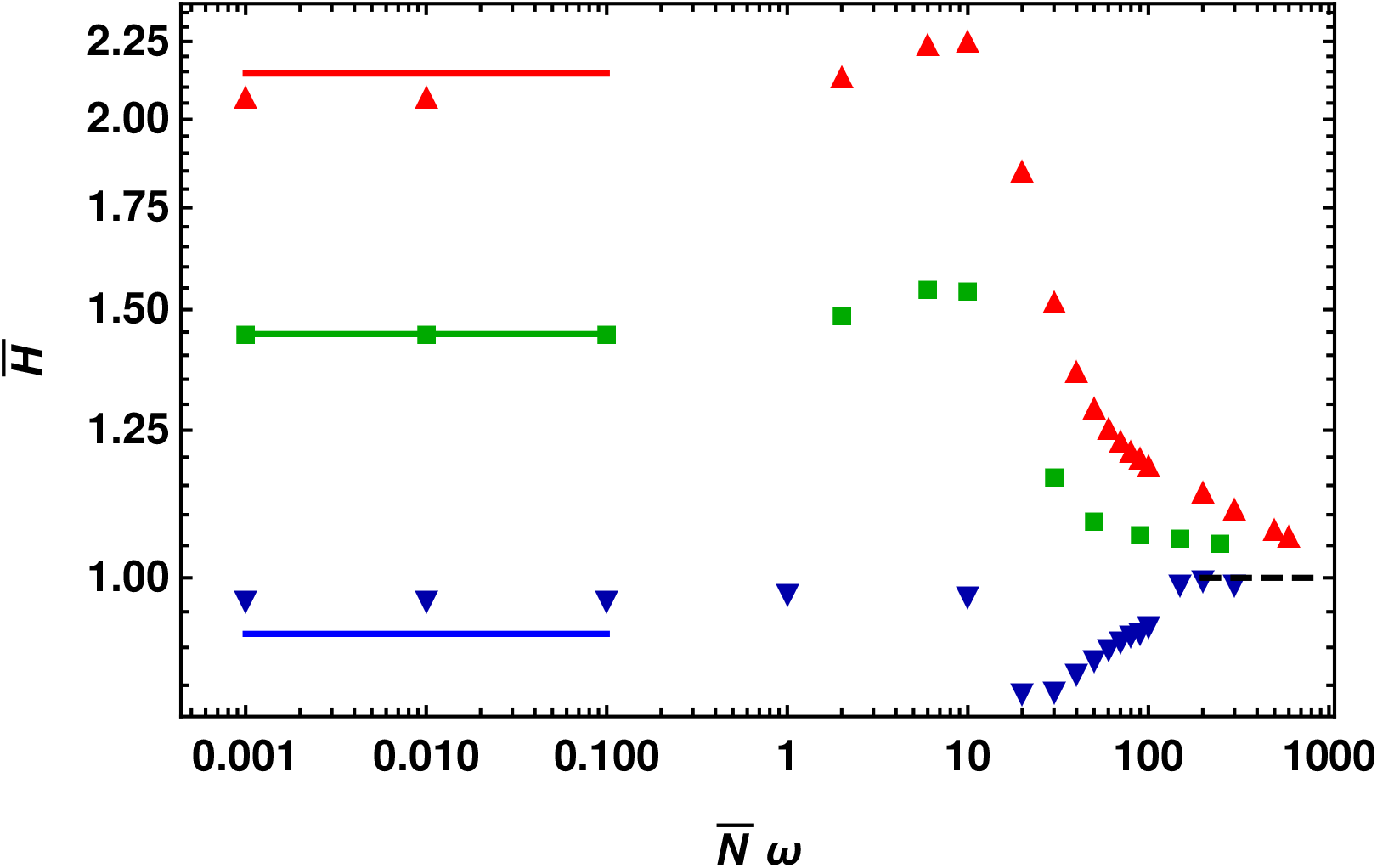
Changing environment and on-average neutral selection: The time-averaged mean heterozygosity as a function of scaled environmental frequency for the dominance parameter *h* = 0.3(▼), 0.5(■) and 0.7(▲) when both selection coefficient and population size change in time with equal frequency. The points are obtained by numerically solving (5) and (9), and the solid lines on the left show the analytical approximation (30) for slowly changing environment while the dashed line on the right is the equilibrium mean heterozygosity *θ* in the time-averaged environment. The other parameters are *ν* = 0.7, *θ* = 1, and 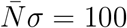. For comparison, note that the mean heterozygosity in neutral constant environment is equal to *θ* = 1.

Our key results summarized in Fig. 5 for on-average neutral selection and various environmental scenarios suggest that in addition to demography, varying selective environment (about zero mean) is a potentially important factor for explaining the lower levels of neutral genetic diversity than predicted on the basis of constant neutral population assumption (for a recent overview, see Buffalo (2021)). In a recent study, we have shown that changing selective environment breaks the symmetry between the conditional fixation times for beneficial and deleterious mutations which leads to different levels of neutral heterozygosity at a linked site due to beneficial and deleterious sweeps, and that the neutral heterozygosity is strongly affected due to the deleterious sweeps even when the selection is changing slowly while the beneficial sweeps are not much impacted (Kaushik and Jain, 2021). In this article, the fundamental quantity of interest is the mean absorption time and the key effect is that as a newborn mutant is more likely to be lost when selection is negative than when it is positive, and in a slowly changing environment, the contribution to polymorphism will come only from the part of the cycle when the selection is strongly positive, the time-averaged quantities will be quantitatively different from those due to positively selected mutants in constant environment. Periodic inflation and contraction of population size can further affect the genetic diversity.

It is thus clear that even slowly changing environments can produce different qualitative patterns and quantitatively different levels of genetic diversity, and detailed investigations of the joint effect of changing selective environment and demography in more complex scenarios such as when genetic hitchhiking occurs (Barton, 2000) remains a task for the future.

## Acknowledgment

We thank Aurélien Tellier for helpful discussions.

## Appendix

## Appendix A

### Eigenfunction expansion of the site frequency spectrum

The density *g*(*x, t*) obeys (6) with homogeneous boundary condition *g*(1, *t*) = 0 and inhomogeneous boundary condition *g*(0, *t*) = *θρ*(*t*). But one can shift *g*(*x, t*) using a (non-unique) simple function that allows one to work with homogeneous boundary conditions (see, for example, Chapter 8 of Mathews and Walker (1970)). We therefore write

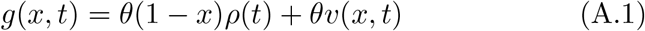

where *v*(*x, t*) satisfies an *inhomogeneous* partial differential equation,

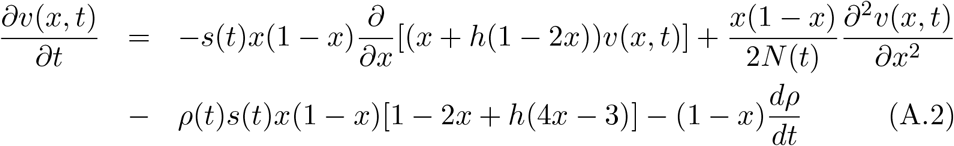

with *homogeneous* boundary conditions, *v*(0, *t*) = *v*(1, *t*) = 0 and initial condition *v*(*x*, 0) defined through (A.1).

The homogeneous boundary conditions allows one to use the method of separation of variables, and therefore one can expand *v*(*x, t*) as a linear combination of the eigenfunction *ψ*_*n*_(*x*) of the homogeneous operator on the RHS of (A.2) with constant selection and fixed population size. Unfortunately, the eigenfunctions *ψ*_*n*_(*x*) are not known in a closed form when selection is present (Kimura, 1957), and we will therefore study (A.2) in Appendix C in the absence of selection.

## Appendix B

### Equilibrium site frequency spectrum for strong selection

In order to obtain an approximate expression for the SFS when the environment is static and selection is strong, we first note that

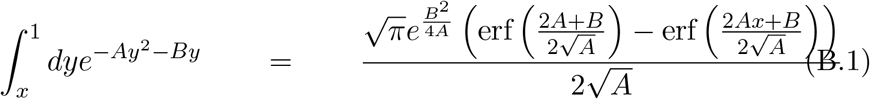

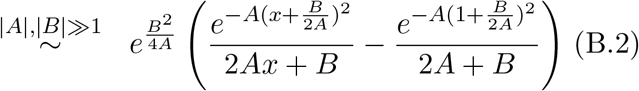

where the last expression is obtained using the asymptotic expansion of the error function for large argument (see (7.1.23) of Abramowitz and Stegun (1964)), and holds provided 2*Ax* + *B* and 2*A* + *B* are nonzero.

Using this result in (11), we find that in constant environment, when the population is under strong selection 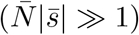 and *h* ≠ 0, 1, the stationary state SFS can be approximated as

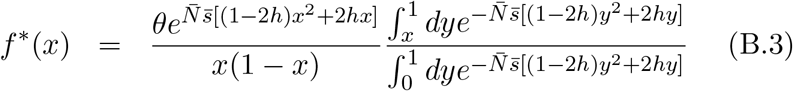

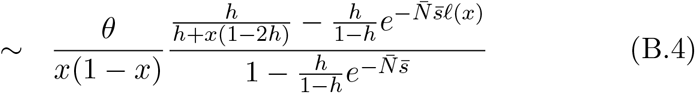

which further leads to

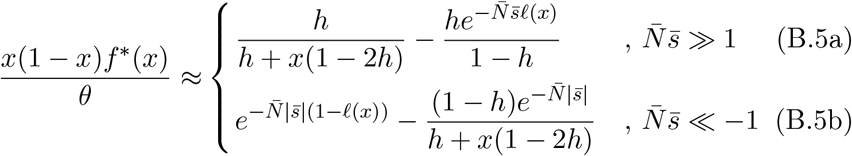

where *ℓ*(*x*) = 1 − *x*^2^ − 2*h*(*x* − *x*^2^) = (1 − *x*)[(1 − *x*) + 2*x*(1 − *h*)] lies between zero and one. If *x* is not close to one, the subleading (second) term in the above equations can be neglected, and we arrive at (13a) and (13b) in the main text.

Using (B.5a) and (B.5b) in (8) for the sample SFS, we obtain

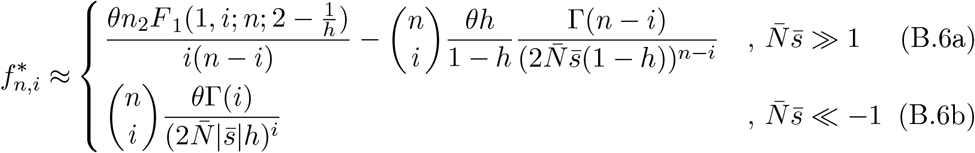

where Γ(*i*) is the gamma function and _2_*F*_1_(*a, b*; *c*; *z*) is the Gauss hypergeometric function (Abramowitz and Stegun, 1964). To obtain the above result for negative selection, we needed to evaluate 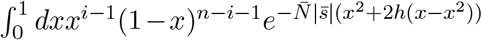, which may be approximated by 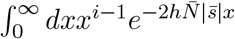 on noting that the main contribution to the integrand comes from small *x* when 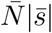 is large. For positive selection, the second term on the RHS of (B.6a) is obtained in a similar fashion, and shows how the asymptotic result is approached.

We briefly discuss the equilibrium SFS for a completely recessive and dominant allele. For *h* = 0, we can use (B.2) for the integral in the numerator of the RHS of (B.3). But as 2*Ax* + *B* = 0 for *x* = 0, we use the asymptotic expansion of error function for the integral in the denominator, and finally obtain

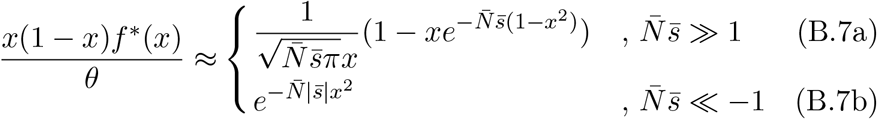

Similarly, for *h* = 1 and 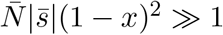, we get

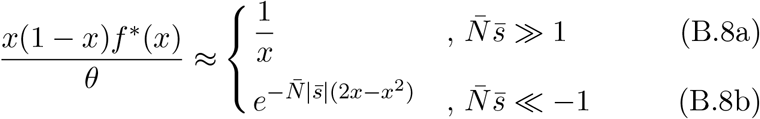

## Appendix C

### Nonequilibrium site frequency spectrum for neutral population

Although the moments of *g*(*x, t*) have been studied when selection is absent and the population size is an arbitrary function of time (Evans *et al*., 2007; ŽivkoviĆ and Stephan, 2011), an explicit expression for *g*(*x, t*) for periodically changing population size has not been obtained. For *s*(*t*) = 0 at all times, from (A.2), we have

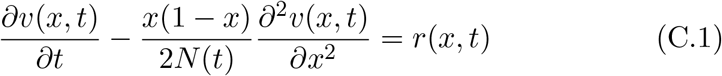

where 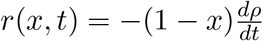. We first write

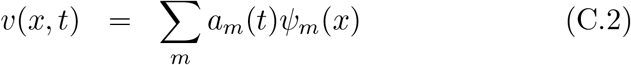

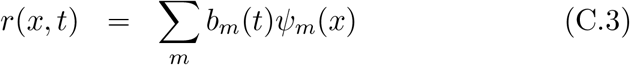

where the eigenfunction *ψ*_*n*_ obeys the eigenvalue equation 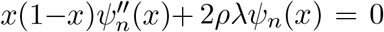 with boundary conditions *ψ*_*n*_(0) = *ψ*_*n*_(1) = 0, and is orthonormal with respect to the weight function *w*(*x*) = [*x*(1 − *x*)]^−1^, that is, 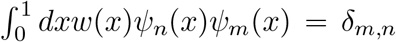. It is known that the eigen-function 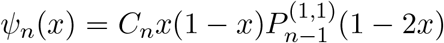 with the eigenvalue *λ*_*n*_ = *n*(*n* + 1)/(2*N*), *n* = 1, 2, … (Kimura, 1964) where 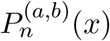 is the Jacobi polynomial and the normalization constant *C*_*n*_ is given by (see (22.2.1) of Abramowitz and Stegun (1964))

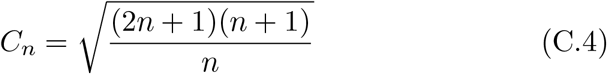

Substituting (C.2) and (C.3) in (C.1) and using the orthonormality condition for the eigenfunctions, we find that the time-dependent coefficient *a*_*m*_(*t*) is the solution of the following first order ordinary differential equation,

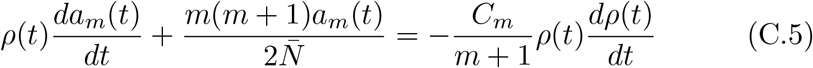

Putting the above discussion together in (A.1), we find that the site frequency spectrum for a large population is given by

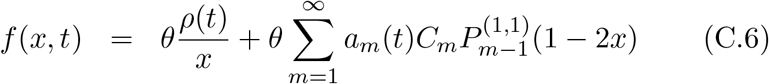

Furthermore, using the above expression and (7.391.2) of Gradshteyn and Ryzhik (2007), the sample SFS is found to be

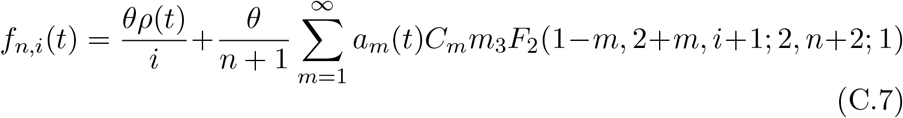

where _3_*F*_2_(*a*_1_, *a*_2_, *a*_3_; *b*_1_, *b*_2_; *c*) is the generalized hypergeometric function. Using the moments of *g*(*x, t*), the above result has also been obtained by ŽivkoviĆ and Stephan (2011).

For the periodically varying population size, the homogeneous solution of (C.5) decays to zero at large times and the inhomogeneous solution is a periodic function with frequency Ω. Therefore, at large times, the time-dependent part *a*_*m*_(*t*) can be expanded in a Fourier series as

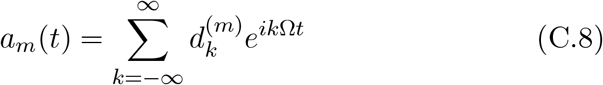

Substituting the above equation in (C.5), we find that the coefficients 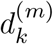’s obey the following three-term recursion equation,

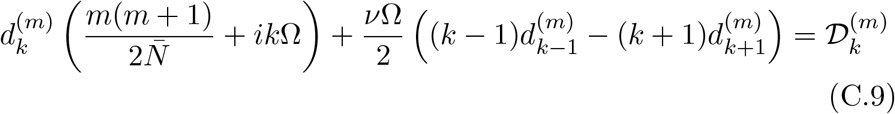

with 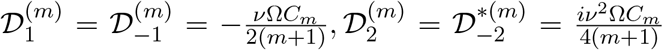 and 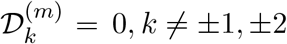 where the subscript * denotes the complex conjugate. It is clear from (C.9) that 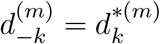 which ensures that *a*_*m*_(*t*) is real.

The above recursion equation does not seem to be exactly solvable but we can obtain approximate expressions for the 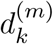’s when the cycling frequency is small or large relative to the inverse average population size.

#### Small cycling frequencies

We first consider the parameter regime 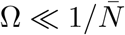. On adding (C.9) for *k* = ±1, we obtain

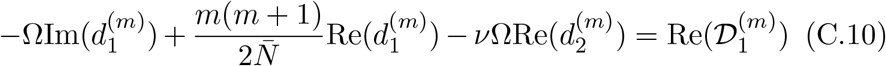

Since *a*_*m*_(*t*) and hence 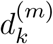’s must vanish with Ω, it follows that the first and third term on the LHS of the above equation are of order Ω^2^ or higher. But since the RHS is of order Ω, we have 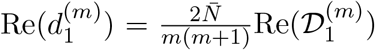. Using this result in (C.9) for *k* = 0, we find that

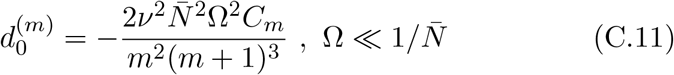

The higher coefficients can also be obtained, and to order 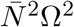, we get

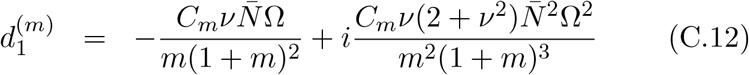

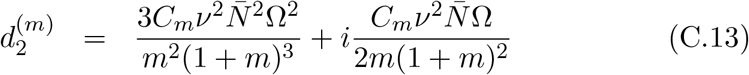

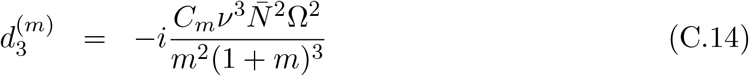

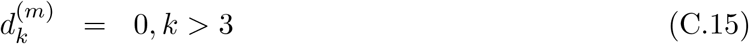

From (C.7), the time-averaged sample SFS can be written as

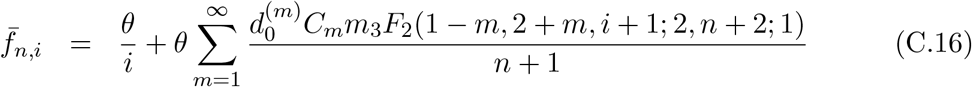

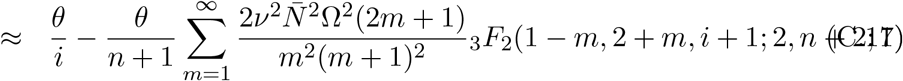

where the last expression is valid for small cycling frequencies. The time-averaged mean heterozygosity can be obtained from the above equation on using _3_*F*_2_(1 − *m*, 2 + *m*, 2; 2, 4; 1) = *δ*_*m*,1_ and given by

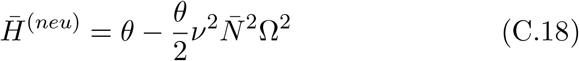

#### Large cycling frequencies

For 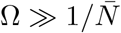, the recursion equation (C.9) for *k* ≠ 0, ±1, ±2 can be approximated by 2*ix*_*k*_ + *ν*(*x*_*k*−1_ − *x*_*k*+1_) ≈ 0 where 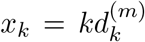. The solution of this equation is given by 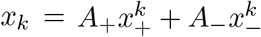 where

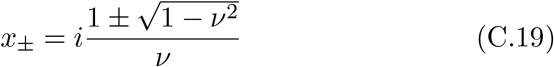

In order that the Fourier series (C.8) converges, for positive *k*, we demand 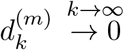 which yields the coefficient *A*_+_ = 0. Using the solution for 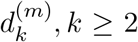 in (C.9) for *k* = 1, 2, we can find *A*_−_ and *d*_1_ to leading order in 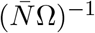, and finally obtain

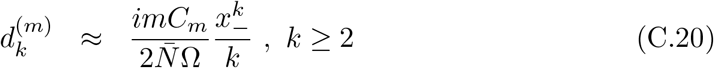

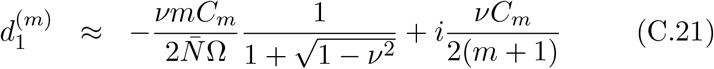

Using the above result for 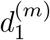 in (C.9) for *k* = 0, we get

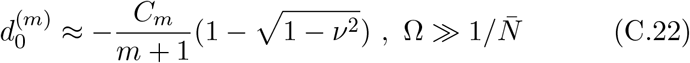

Thus, for large cycling frequencies, the time-averaged sample SFS is given by

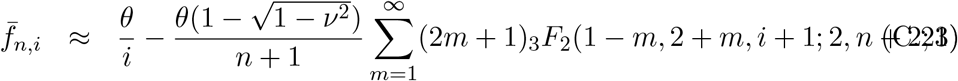

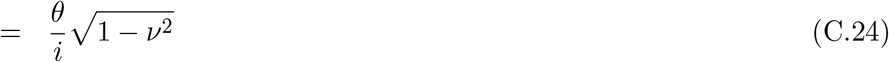

where the last expression follows on using that the sum on the RHS of (C.23) equals (*n* + 1)/*i*, as can be verified numerically.

## Appendix D

### Nonequilibrium site frequency spectrum for weak selection

Here we develop a perturbation theory for weak selection when both the selection and population size oscillate with the same frequency. In the following discussion, we will write *s*(*t*) = *♯ζ*(*t*) where 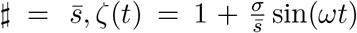 and *♯* = *σ, ζ*(*t*) = sin(*ωt*) when on-average selection is nonzero and zero, respectively. We begin by writing 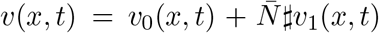 and substitute it in (A.2). Keeping terms up to linear order in 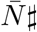, we find that *v*_0_ obeys (C.1) and *v*_1_ is a solution of the following equation,

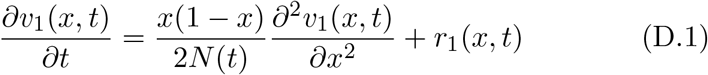

where,

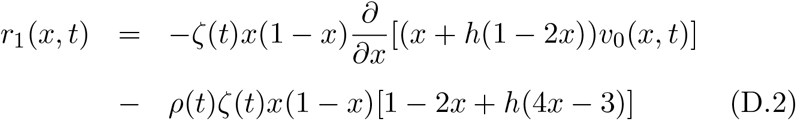

As in Appendix C, we expand *v*_1_ and *r*_1_ as a linear combination of the eigenfunctions *ψ*_*m*_(*x*),

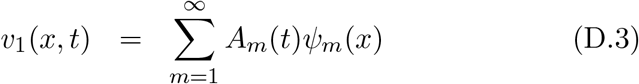

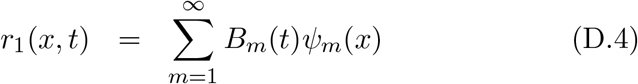

where *A*_*m*_(*t*) obeys the following ordinary differential equation,

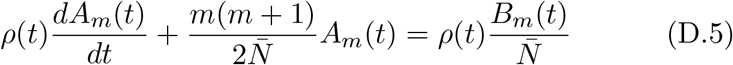

and its long time solution can be expanded in a Fourier series as 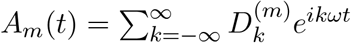. The coefficient 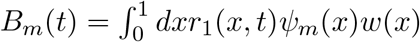 can be found using the orthonormality condition for the eigenfunctions (see Appendix C). But as the integrals involved in *B*_*m*_(*t*) do not seem to be known for arbitrary *m*, here we will focus on *B*_1_(*t*) which is related to mean heterozygosity and given by

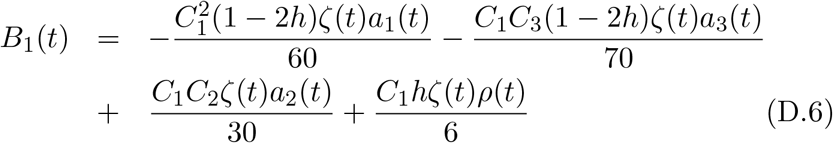

where *a*_*m*_(*t*) is defined in (C.8). The sample SFS averaged over a period can be written as

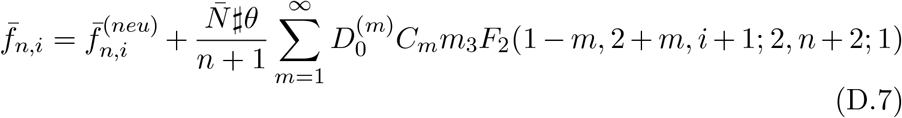

where 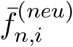 is given by (C.17). In particular, for the time-averaged heterozygosity, we have

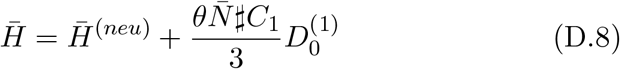

where 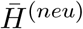 is given by (C.18) for small cycling frequencies and 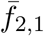 in (C.24) for large cycling frequencies.

To demonstrate the weak selection calculation, below we consider the case when 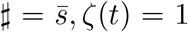, that is, selection is constant. We find that the coefficients 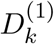 obey the following recursion equation,

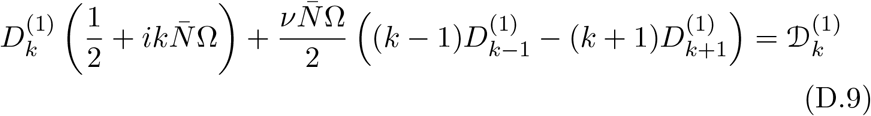

where

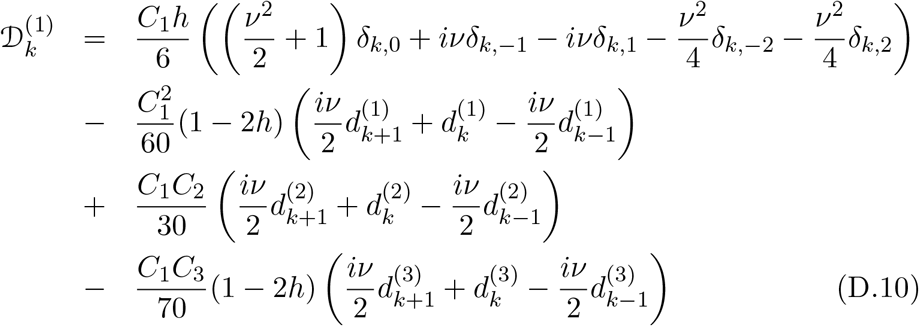

For small cycling frequencies, using an argument similar to that given in Appendix C for 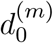, we get 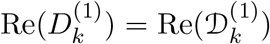; on using 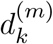 given by (C.11)-(C.15), we find that 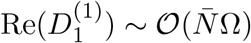. But as 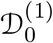 is a constant in 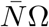, we finally obtain

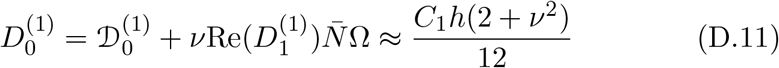

For large cycling frequencies, as in Appendix C, 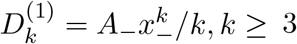. Then using the properties of 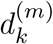 for |*k*| < 3 described above, we find

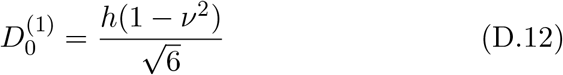

## Appendix E

### Mean heterozygosity for co-dominant, on-average neutral mutant

For co-dominant allele in an on-average neutral environment with constant population size, the time-averaged mean heterozygosity is equal to *θ* for any cycling frequency and strength of selection. For 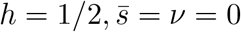, (6) reduces to

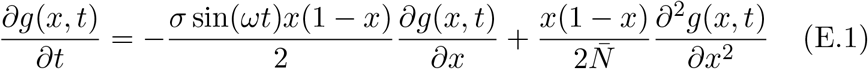

We first note that the above equation has the following symmetry:

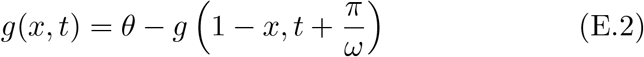

where the boundary conditions, *g*(0, *t*) = *θ, g*(1, *t*) = 0 have been used. Then using the definition 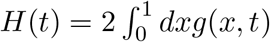 and on integrating by parts, (E.1) yields

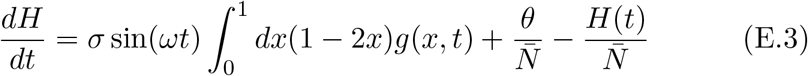

Integrating both sides over the period 2*π*/*ω*, we find that the LHS vanishes since *H*(*t*) is a periodic function, and the first term on the RHS is also zero due to the symmetry property (E.2); we thus finally arrive at 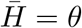.

## Appendix F

### Nonequilibrium site frequency spectrum for strong selection

For on-average neutral selection and equal cycling frequencies for selection and population size, we write (6) as

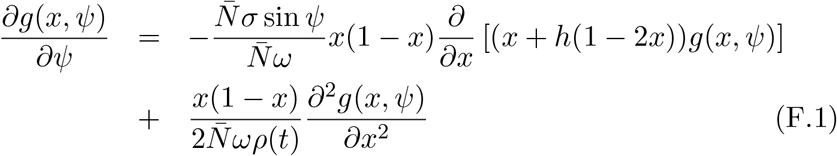

where *ψ* = *ωt* and expand *g*(*x, t*) in powers of 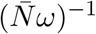. If 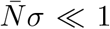 and 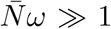, the first term on the RHS must be neglected in comparison to the second term on the RHS, and therefore the leading order result for *g*(*x, t*) is given by the neutral population already discussed in Appendix C and the subleading correction due to selection in Appendix D. For strong selection, if the LHS is also ignored, we get *g*(*x, t*) is a constant in allele frequency which is not true.

Here, we are therefore interested in the regime where 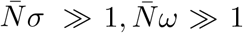 with *σ*/*ω* finite. Then for rapidly changing environment and strong selection, the second term on the RHS can be ignored resulting in a first order partial differential equation for *g*(*x, t*). A numerical analysis of the resulting approximate equation with boundary conditions (7) is found to be in reasonable agreement with that for the exact equation. For simplicity, we consider the case when *h* = 1/2 and solve the first order partial differential equation by the method of characteristics.

For this purpose, we consider a change of variables as *ξ* = *ξ*(*x, ψ*) and *η* = *η*(*x, ψ*), and choose *ξ* = *ψ* and *η* = constant so that

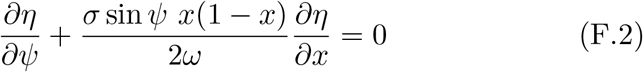

which leads to the characteristic equation,

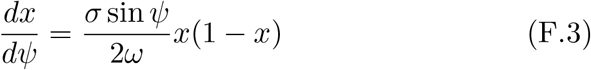

and the characteristic curve to be,

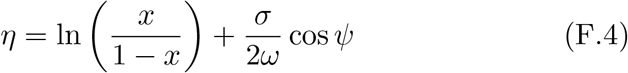

We then obtain *∂g*/*∂ξ* = 0 which implies that *g*(*ξ, η*) = 𝒢(*η*) where 𝒢 should be found using the boundary conditions. We are, however, unable to implement the boundary conditions (7) in this solution. If instead we determine 𝒢 using the initial condition, *g*(*x*, 0) = *θ*(1 − *x*), we get

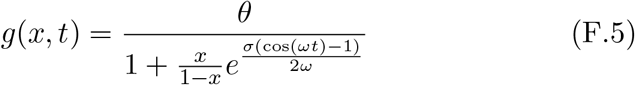

which shows that the *g*(*x, t*) has phase difference of *π*/2 from selection, but this result is independent of *ν* and Ω which is inconsistent with the numerical results for *g*(*x, t*).

## Notes

### Competing Interest Statement

The authors have declared no competing interest.

